# Inter-hemispheric inhibition in stroke survivors is related to fatigue and cortical excitability

**DOI:** 10.1101/831511

**Authors:** Sasha Ondobaka, Nick Ward, Annapoorna Kuppuswamy

## Abstract

**Objective:** Persistent post-stroke fatigue is a major debilitating condition that has been linked to low corticomotor excitability and aberrant attention, both phenomena that are associated with the inter-hemispheric inhibition balance in the brain. In this study, we examined the relationship between inter-hemispheric inhibitory effective connectivity, motor cortex excitability and chronic persistence of post-stroke fatigue.

**Methods:** We tested eighteen non-depressed stroke survivors with minimal motoric and cognitive impairments using spectral dynamic causal modelling (spDCM) of ‘resting state’ magnetic resonance imaging (rs-fMRI) and transcranial magnetic stimulation (TMS) measures of cortical excitability. We also assessed the levels of non-exercise induced, persistent fatigue using Fatigue Severity Scale (FSS) - a self-report questionnaire which has been widely applied and validated across different conditions. To understand neural effective connectivity mechanisms involved in fatigue and corticomotor excitability we examined the balance in inhibitory connectivity between homologue regions in M1, anterior insula, caudate and thalamus of the resting brain.

**Results:** Inter-hemispheric inhibition balance between left and right M1 accounted for 67% of variability in the reported fatigue (R=.82, p<0.001). Inter-hemispheric inhibition balance in M1 also accounted for 54% of variability in the corticomotor excitability characterised by individual resting motor thresholds (R=.74, p<0.001), a measure that has been associated with subjective fatigue reports. Other examined inter-hemispheric connections did not show significant relationships with either fatigue or cortical excitability measures.

**Conclusion:** Our findings suggest that the balance in inter-hemispheric effective connectivity between primary motor regions is involved in regulation of corticomotor excitability and could explain subjective post-stroke fatigue.

## INTRODUCTION

Fatigue is a major debilitating symptom in many neurological and psychiatric disorders [1], including stroke. The reported prevalence of fatigue in stroke survivors is as high as 85%, with post-stroke fatigue having a significant impact on stroke survivors’ disability, quality of life and mortality [2–4]. Both central and peripheral mechanisms have been implicated in the development and persistence of post-stroke fatigue [5–7]. Peripherally, high levels of tissue inflammation early after stroke is linked with a subsequent genesis of post-stroke fatigue [8–10]. Centrally, persistent post-stoke fatigue is associated with low cortical excitability and aberrant neural regulation of attention [11,12], yet the underlying neural mechanisms remain unknown.

Inter-hemispheric inhibition balance (IIB) has been proposed as a neurophysiological mechanism that could underlie both attention and cortical excitability. In the normal functioning brain, inter-hemispheric connectivity is not balanced between hemispheres, but exhibits an asymmetry characterised by a net left to right inhibitory dominance in the motor cortices [13–15], and outside primary motor areas [16,17]. In this study, we hypothesised that the directionality and the extent of inter-hemispheric inhibition balance could provide a mechanistic explanation for inter-individual variability in post-stroke fatigue, which could serve as a therapeutic target in the future.

To test this hypothesis, we applied a computational modelling approach to resting state functional MRI connectivity. We predicted that the deviation from left to right inter-hemispheric dominance would be positively associated with both severity of persistent self-reported post-stroke fatigue and the associated cortico-spinal excitability [13–15,17]. Specifically, we predicted that participants with low fatigue would show left inhibitory inter-hemispheric dominance, whereas those with high fatigue would show more symmetrical IIB or tend towards right inhibitory dominance.

## METHODS

### Participants

The sample consisted of 18 (two female) right-handed non-depressed, ischemic or haemorrhagic stroke survivors with a mean age of 58.67±10.61 (mean ± SD), tested 27.71 [35.90] (median [inter-quartile range]) months post stroke. The study was approved by the Riverside Research Ethics Committee (12/LO/1474).

#### Inclusion criteria

The diagnostic inclusion criteria for recruitment included a clinical diagnosis of a first-time ischaemic or haemorrhagic lesion, survivors with no subarachnoid haemorrhage and transient ischaemic stroke.

#### Exclusion criteria

To avoid potential sources of bias, following exclusion criteria were adopted: use of centrally acting medication, high score on the Hospital Anxiety and Depression Scale (>11) and poor function. Functional screening included upper limb functional tests and cognitive tests. Poor upper limb function was defined as having <60% of the unaffected limb score in more than one of the following measures: (i) Nine Hole Peg Test (9HPT) to measure finger dexterity; (ii) Action Research Arm Test (ARAT); (iii) Grip strength. Poor cognitive function was defined as a score 45 on the Sustained Attention Index (SAI) and Symbol Digit Modalities Test (SDMT) as a measure of information processing speed.

#### Participant recruitment

During a period between February 2013 and September 2014, a total of 225 stroke survivors were consecutively screened for compatibility with TMS and fMRI procedures. The screening was done via the Thames Stroke Research Network from the University College NHS Trust Hospital, Epsom NHS Trust Hospital, Royal Surrey NHS Trust Hospital and from community stroke groups. Eligibility criteria were met by 78 participants, of which we have recruited 70 for the TMS session (reported in Kuppuswamy et al., 2015). 18 of the 70 stroke survivors additionally took part in the resting state fMRI scanning session and were included in this study and their data was used for the reported analyses. There were two participants with missing data for the SAI measurement and one participant with a missing date of birth.

### Measurements

We assessed the level of non-exercise induced, persistent fatigue using Fatigue Severity Scale (FSS) - a validated self-report questionnaire, commonly used in various different conditions [18]. In two experimental sessions, we measured low-frequency spontaneous fluctuations in the resting state functional magnetic resonance imaging (rs-fMRI) signal and resting motor thresholds (RMTs), a typical measure of corticospinal excitability. Our use of RMTs was motivated by its association with fatigue in our previous work. For full details of the TMS procedure used in the first session please refer to our previous publication [19]. In the second session, participants underwent an eyes-open twelve-minute rs-fMRI using a standard scanning protocol. Scanning was performed at the Wellcome Centre for Human Neuroimaging using a 3T Trio scanner (Erlangen, Siemens). All participants underwent a single scanning session during which a T2*-weighted MRI transverse echo-planar images (EPI) were acquired using a 12-channel head coil. The resting block comprised 200 volumes of 32 slices, with a 30ms echo time (TE) and a repetition time (TR) of 2.175 s (4.5 × 4.5 × 4.5 mm voxels). Participants were instructed to lie within the scanner, to keep their eyes closed, to remain awake and to restrict their movement as much as possible until further instructed. A high resolution T1-weighted anatomical images (1.3 × 1.3 × 1.3 mm voxels; 176 partitions, FoV = 256 × 240, TE = 2.48ms, TR = 7.92ms, FA = 16°) and a field map (TE1 = 10ms and TE2 = 12.46ms, 3×3×3mm resolution, 1mm gap) were also acquired. This was used to create B0 fieldmaps used by the SPM Fieldmap Toolbox to unwarp the functional images. Raw data can be made available upon request.

### fMRI pre-processing

We performed conventional functional imaging pre-processing using SPM12 (www.fil.ion.ucl.ac.uk/spm), including the removal of the first four volumes, realignment, spatial normalization with 3mm cubic voxels, a spatial smoothing of 6 mm FWHM and nuisance variable regression. The nuisance regressors included 18 motion parameters (six head motion parameters and their first and second derivatives) and the average signal strength extracted from 6mm spheres from two (cerebrospinal fluid and white matter) reference regions with the following MNI coordinates: 19 −34 18 and 27 −18 32. This set of nuisance variables incidentally removes low-frequency fluctuations normally associated with global confounds.

### Dynamic causal modelling of neural effective connectivity

SpDCM provides a model-based approach to characterise biologically plausible effective neural connectivity mechanisms that cause changes in the measured blood-oxygen-level dependent (BOLD) signals. Compared to the classic functional connectivity methods that assess correlations between BOLD signal intensity in different regions [20,21], spDCM offers a possibility to discern the directionality and the sign (net inhibition vs excitation) of the underlying neural influences [22,23]. spDCM models extrinsic (i.e. between regions) and intrinsic (i.e. within region) neural effective connectivity in the selected brain regions of interest based on the estimated cross-spectra (cross-covariances in the frequency domain) of the extracted BOLD time-series. An advantage of using a model-based approach like DCM is the opportunity to infer the biologically plausible inter-hemispheric neural dynamics that underlie fatigue, as compared to investigating the dynamics in the measured EEG or fMRI signals. Examination of the model parameter values that provide best fit for the observed fMRI signals allow characterisation of the biologically plausible directionality and the sign of the inter-hemispheric connectivity patterns that best explain reported fatigue and measured corticospinal excitability.

To employ spDCM we extracted rs-fMRI BOLD time-series from bilateral primary motor cortices (M1; 30 −24 64; −30 −20 66), anterior insular cortices (38 16 2; −36 16 0), thalamus (12 −12 10; −12 −12 10) and caudate nuclei heads (14 12 14; −14 12 14). Additionally, we included the three cortical midline regions of interest (ROIs): the supplementary motor area (SMA; 0 - 6 58) and the two key default mode regions, ventromedial prefrontal cortex (vmPFC; 0 50 −4) and posterior cingulate cortex (PCC; 0 −52, 24). Exact locations of the centres of the 8mm ROI spheres, indicated above in MNI coordinates, were determined by using NeuroSynth meta-analysis maps. NeuroSynth (http://neurosynth.org) is an online platform for large-scale automated meta-analysis of published neuroimaging results that provides posterior probabilities of a specific term being used in the abstract of the analysed publications (e.g. primary motor cortex) conditional on the presence of activation in a chosen voxel. We selected the voxels with the highest posterior probability of being associated with the terms of the 11 chosen ROIs included to capture the global neural dynamics of both hemispheres (**Figure 1**). Visual inspection of structural images showed no apparent lesions in any of the selected ROIs. The resulting specified model consisted of a fully connected architecture with 110 extrinsic connections and 11 intrinsic connections (**Figure 1**).

**Figure 1.**
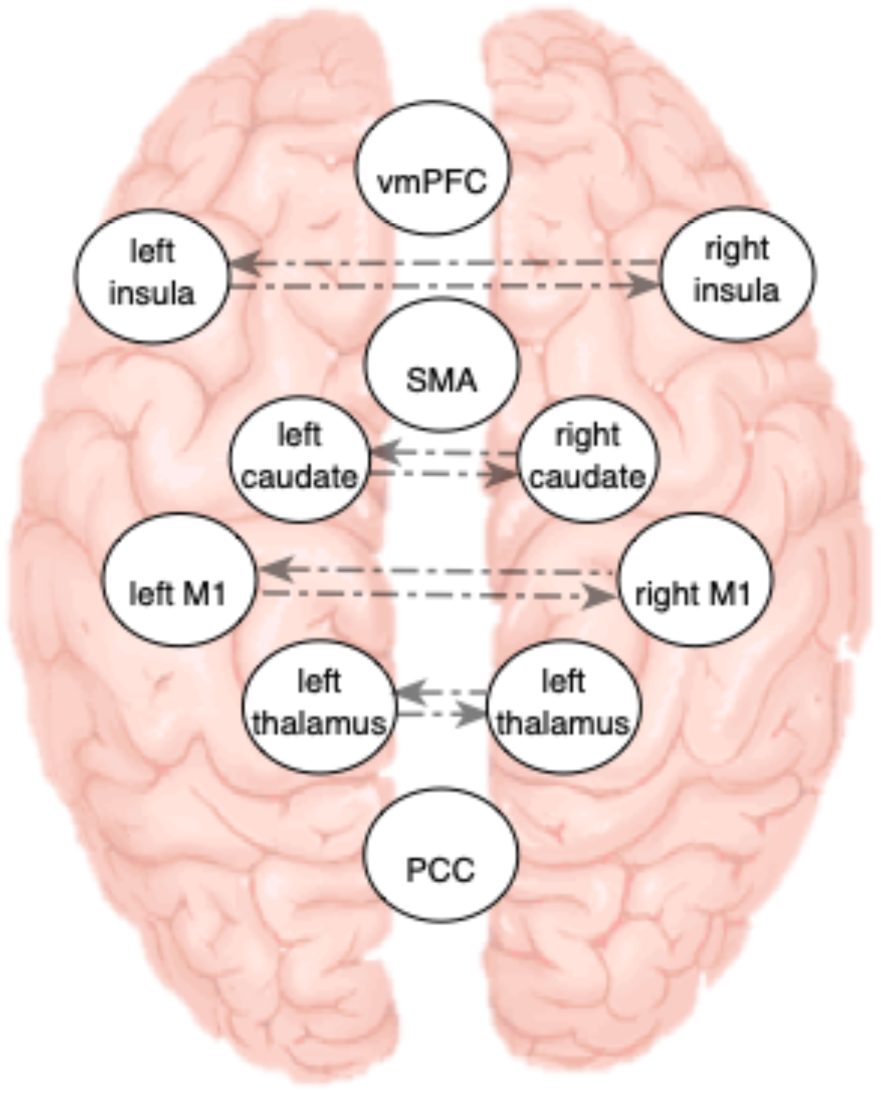
DCM architecture. This figure represents the dynamic causal model architecture consisting of 11 brain regions selected from NeuroSynth tool (http://neurosynth.org/; ventromedial prefrontal cortex (vmPFC), left and right anterior insula, supplementary motor area (SMA), left and right caudate head, left and right primary motor cortex (M1), left and right thalamus and posterior cingulate cortex (PCC)). The model was fully connected, consisting of 110 extrinsic (between regions) influences, and 11 intrinsic (within region) influences. Depicted are the 8 inter-hemispheric influences, other connections are left out from the figure for clarity. We used this biologically plausible dynamic causal model of intrinsic and extrinsic neuronal connectivity to find the best fit or explain the recorded fluctuations in BOLD intensities.

### Participants-specific Bayesian estimation of effective connectivity parameters

At the first, participants-specific level of analysis, we estimated the strength of all effective connectivity parameters for each participant using a standard Bayesian inversion scheme (Variational Laplace; [24]). During the iterative estimation (model inversion) procedure, the connectivity parameter strengths representing biologically plausible extrinsic and intrinsic influences are fitted to the BOLD data using Bayesian inference. DCM uses negative variational free energy to approximate the log-evidence that a particular model of connectivity patterns fits or explains the observed dynamics in the region-specific BOLD time-series.

To account for random effects on connectivity from participants to participants, we used a hierarchical parametric empirical (PEB) Bayesian model that furnishes a more efficient and robust estimation of effective connectivity parameters [25]. The participants specific estimates were used for subsequent analysis. In DCM, estimated positive values of extrinsic parameters represent excitatory influence and negative values index inhibitory influence from region A to region B, while their absolute values represent percent estimated activity change of this influence (effect size).

### Inter-hemispheric inhibition balance (IIB) index of effective connectivity

Using participants specific posterior expectations for each effective connection, we computed the inter-hemispheric inhibitory balance (IIB) index for primary motor cortex, the anterior insular cortex, caudate and the thalamus by subtracting right-to-left hemisphere parameter values from left-to-right parameter values [IIB = (L to R) - (R to L)]. IIB index characterises the nature of normally occurring inter-hemispheric inhibitory balance in the individual ‘resting’ brain. Negative IIB values reflect stronger left to right net inhibitory influences, whereas positive values reflect stronger inhibitory right to left influences. To better explain endogenous neuronal fluctuations and get a better estimate of IIB, in addition to the motor regions, we also considered key default mode network nodes (posterior cingulate cortex and ventro-medial prefrontal cortex), and the supplementary motor area.

### Group-level multiple regression of FSS and RMT values from the inhibitory balance (IIB) indices

Finally, at the between-participants (group) level analysis, we used the participants-specific IIB indices and the interaction between paretic side (left vs right) and IIB indices as independent explanatory variables of: 1) individual differences in subjectively reported FSS (fatigue) scores, and 2) individual resting motor thresholds (RMTs) measured by MEP amplitudes. The interaction term was included to test for a potential confounding effect of paresis. We used classic multiple linear regression analysis in MATLAB 2006b (The MathWorks, Inc., Natick, Massachusetts, United States) to separately test whether the explanatory variables significantly explain variance in the two target variables. We applied Bonferroni correction to a chosen significance threshold of *p* < 0.05 to control for false positives over four tests of inter-hemispheric inhibitory balance in M1, insula, caudate and the thalamus, separately for the target variables FSS and RMT. To better understand the nature of IIB, we have also examined the individual left to right and right to left connections between the four homologous regions. We furthermore tested for the statistical dependence between the individual FSS and RMT values in our sample by calculating the Pearson’s correlation coefficient.

## RESULTS

### Participants characteristics

Seven participants had a paresis on their left side, 11 on the right - all participants were right-handed. Participant characteristics indicated low cognitive impairment, indexed by unaffected mental speed (SDMT scores of 1.25 ± 0.42) and by low SAI measure of attention 0.44 ± 1.21 (mean ± SD). Participants also showed low motoric impairment, reflected in the measured 9HPT 77.73 ± 33.06 scores, in ARAT scores 96.01 ± 11.17 % of the unaffected hand and preserved grip strength of 88.67 ± 22.14 (mean ± SD) of the unaffected side.

### Relationship between fatigue scores and resting motor thresholds

We observed a significant correlation between the subjectively reported FSS scores and the physiological measures of RMTs (*p* = 0.003, R = 0.443, **Figure 2**). As **Figure 2** shows, the correlation did not depend on the participants’ paretic side that can be taken as a proxy for the presence of the lesion in the contralateral hemisphere.

**Figure 2.**
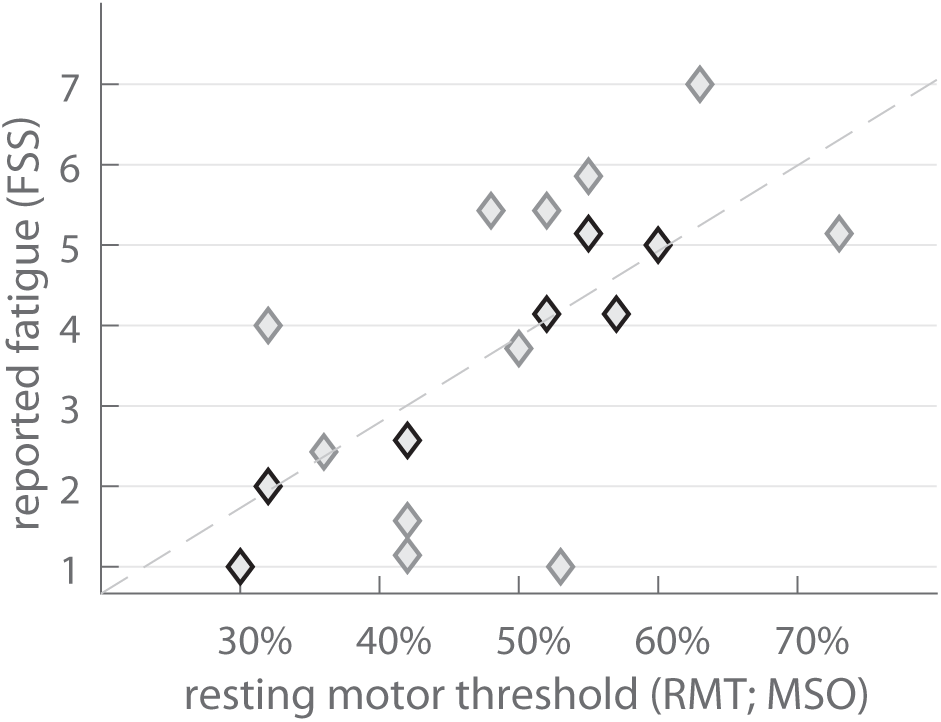
Self-reported fatigue scores (FSS) and resting motor thresholds (RMTs). Graph represents relationship between self-reported FSS values and RMTs measures of corticospinal excitability, indexed by percentage maximum stimulator output (MSO). Grey diamonds mark participants with right side paresis, black diamonds mark participants with left side paresis.

### Inter-hemispheric balance explains reported level of persistent fatigue

The result from the multiple linear regression analysis showed that the individual inter-hemispheric inhibition balance (IIB) in the motor cortex explains reported levels of persistent fatigue. IIB in M1 explained reported FSS scores (t = 5.386, p = 7.56e-05, **Figure 3C)**, whereas Paretic side did not influence the IIB-FSS association (t = 0.112, p = 0.913, for IIB x Paresis interaction), accounting for 67% of the variability in the reported fatigue scores (R = 0.820). After separating the inter-hemispheric neural influences, we observed a positive association between FSS scores and left to right M1 influences (t = 2.636, p = 0.018, **Figure 3A**), such that higher FSS corresponded to stronger excitatory left to right influences. There was a negative association between FSS and the strength of right to left M1 influence (t = −2.730, p = 0.015, **3B**), indicating that higher FSS scores are associated with stronger inhibitory right to left effective connectivity. Paretic side did not affect the association between either left to right M1 influence (t = −0.977, p = 0.344, for Paresis × Influence interaction), nor right to left M1 influence and FSS (t = 0.700, p = 0.494, for Paresis × Influence interaction).

**Figure 3.**
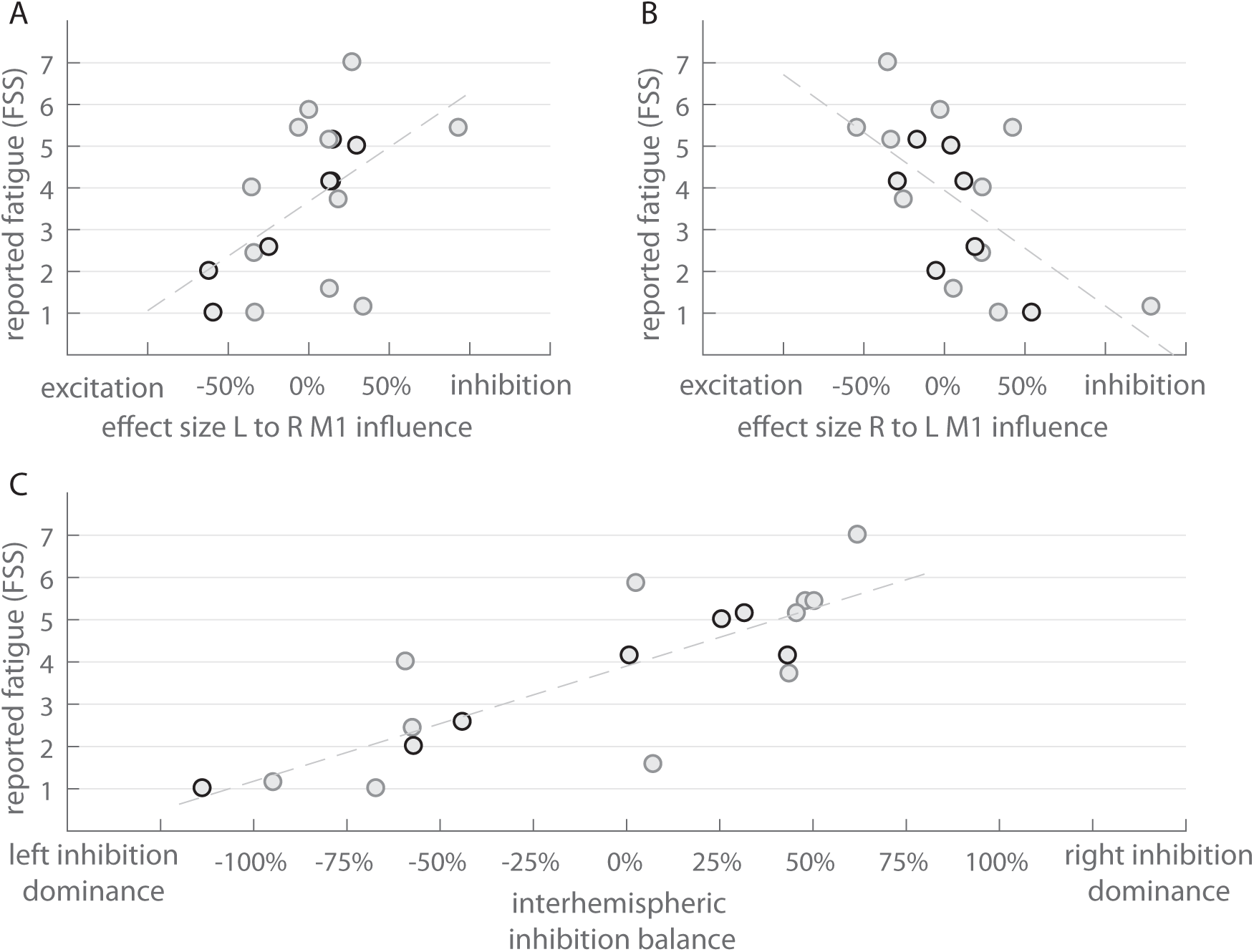
Inter-hemispheric inhibition balance (IIB) in M1 and fatigue severity (FSS). Relationship between the estimated strength of Left (L) to Right (R) M1 influence (effect size, a), and Right to Left M1 influence strength (b) and self-reported FSS. Panel c shows a relationship between IIB indices [L to R minus R to L influence] and self-reported fatigue severity scores. The inter-hemispheric inhibition balance (IIB) was computed by subtracting R to L M1 effect sizes from L to R M1 effect sizes. Negative IIB values on the x-axis reflect overall stronger inhibitory L to R influence, whereas positive values reflect overall stronger inhibitory R to L influence. Strength of effective connectivity is represented by a percentage change in activity (effect size) in an area (e.g. right M1), as a consequence of activity change in another area (i.e. left M1). Grey circles mark participants with the right-side paresis, black circles mark participants with the left-side paresis.

### Inter-hemispheric balance explains measured resting motor thresholds

IIB index in M1 explained the individual resting motor threshold (RMT) values (t = 4.395, p = 5.22e-04, **Figure 4C)**, whereas Paretic side did not influence IIB-RMT the association (t = - 0.876, p = 0.394, for IIB x Paresis interaction), accounting for 56% of the variability in the measured corticospinal excitability (R = 0.750). Again, we observed a positive association between left to right M1 influence and RMTs (t = 3.146, p = 0.007, **Figure 4A**), such that higher RMTs corresponded to stronger excitatory left to right effective connectivity. The strength of right to left M1 connections was negatively associated with RMTs (t = −2.142, p = 0.049, **Figure 4B**), such that higher RMTs corresponded to stronger inhibitory right to left effective connectivity. Paretic side did not affect the association between either left to right M1 influence (t = −1.849, p = 0.084, for Paresis × Influence interaction), nor right to left M1 influence and FSS (t = 0.780, p = 0.447, for Paresis × Influence interaction). We observed no significant effects when the FSS and RMT values were regressed on the IIB scores from the insula, caudate and the thalamus (all *p*s > 0.10).

**Figure 4.**
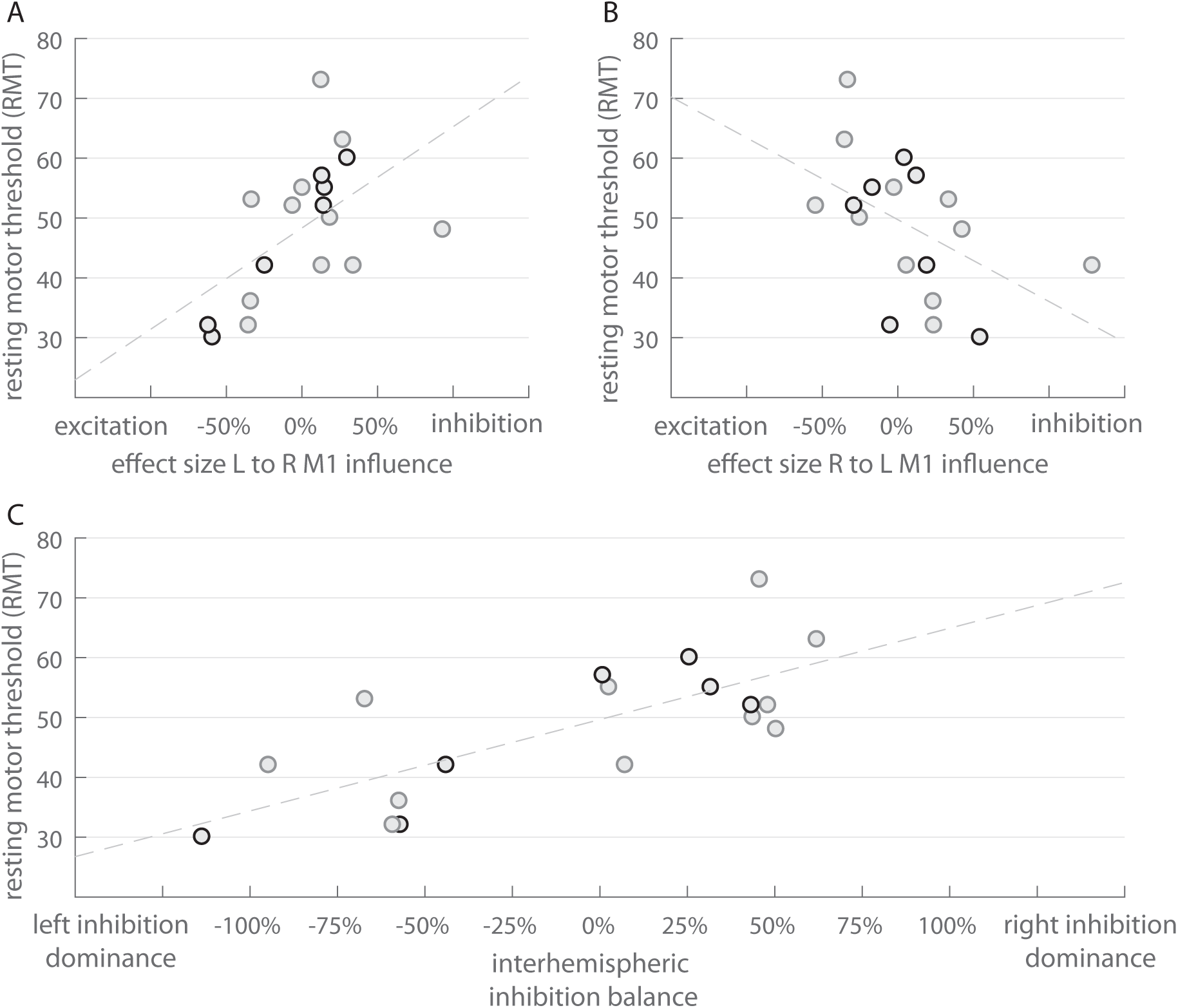
Inter-hemispheric inhibition balance (IIB) and resting motor thresholds (RMTs). Relationship between the estimated Left (L) to Right (R) M1 effective connectivity strength (**a)**, and R to L M1 connectivity strength and measured RMTs (**b**). Panel **c** shows a relationship between IIB indices [L to R minus R to L connectivity] and measured RMTs. The inter-hemispheric balance (IIB) was computed by subtracting R to L M1 strength from L to R M1 strength. Negative IIB values on the x-axis reflect overall stronger L to R inhibitory effective connectivity, whereas positive values reflect overall stronger inhibitory R to L effective connectivity. Strength of effective connectivity is represented by a percentage change in activity (effect size) in an area (e.g. right M1), as a consequence of the activity change in another area (i.e. left M1). Grey circles mark participants with the right-side paresis, black circles mark participants with the left-side paresis.

## DISCUSSION

### Inter-hemispheric balance relates to post-stroke fatigue and level of corticospinal excitability

This study demonstrates that inter-hemispheric network dynamics is altered in post-stroke fatigue and could potentially explain diminished corticospinal excitability, which has previously been related to post-stroke fatigue [19]. Healthy brains at rest exhibit an inter-hemispheric inhibitory balance sustained by the dominant left to right hemispheric inhibition and the complementary right to left hemispheric excitation [13,14]. In our study, opposite patterns of biologically plausible inter-hemispheric inhibitory effective connectivity in primary motor cortices (M1) distinguished stroke survivors with high fatigue from low fatigue and those with high corticospinal excitability from low excitability. The relationship between inferred effective connectivity parameters between left and right M1 and post-stroke fatigue and RMTs was independent of the paretic side, which is taken as a proxy for the hemisphere affected by stroke, indicating that distorted inhibitory balance is unrelated to the location of stroke. Below we discuss our findings in light of prior work and consider potential biological mechanisms that underlie inter-hemispheric connectivity.

### Inter-hemispheric balance (IIB) and cortico-spinal excitability

Our results showed enhanced left to right M1 excitatory influence and right to left inhibitory influence in stroke survivors with relative low corticospinal excitability, and the opposite connectivity pattern in stroke survivors with relative high corticospinal excitability. These findings are in line with data from repetitive transcranial stimulation (rTMS) work showing that inhibitory stimulation to M1 influences cortical excitability in the contralateral M1 [26,27]. Changes in cortical excitability in one M1 elicited by stimulating homologue area in the other hemisphere are dependent on transcallosal connectivity [28]. However, the relationship between altered inter-hemispheric inhibition balance and level of M1 excitability seen here may be mediated by a potential third variable. For instance, both reduction in M1 excitability and inter-hemispheric balance can be viewed as a consequence of the modulatory influence of altered sensory gain (i.e. attention and attenuation) on neuronal connectivity [29]. This view might explain how global changes in sensory attenuation and attention cause observed net inhibitory or excitatory influences as measured by changes in MEP magnitudes or changes in BOLD or EEG signal intensities [30,31].

Change in inter-hemispheric inhibition following stroke, specifically altered inhibitory influence from the unaffected to the affected hemisphere [32–34] resulting in increased corticospinal output, is viewed as a potential mechanism that underlies motor recovery. However, several studies show contradictory results and the contradiction is explained as follows: better post-stroke motor recovery is linked to lower inter-hemispheric inhibition in patients that show greater structural integrity. Conversely, greater inter-hemispheric inhibition is associated with better motor recovery in patients with lesser structural integrity [35]. The seemingly opposite influences from the unaffected hemisphere to increase corticospinal output maybe better explained by taking the combined measure that represents inter-hemispheric balance. The results from the current study suggest that interhemispheric effective connectivity balance as opposed to inter-hemispheric inhibition might be responsible for driving corticospinal output. Perhaps, greater or lesser corticospinal output in relation to motor recovery could also be a result of altered interhemispheric inhibition balance independent of lesion location.

### Inter-hemispheric balance (IIB) and post-stroke fatigue

We related enhanced right hemisphere inter-hemispheric inhibitory dominance to experience of high post-stroke fatigue. Higher left hemisphere inter-hemispheric inhibitory dominance was associated with low post-stroke fatigue. Generally, the opposite pattern of inter-hemispheric excitation-inhibition dynamics in high post-stroke fatigue is in agreement with the disturbed patterns of inter-hemispheric connectivity associated with many neurological and psychiatric disorders [28,36–42]. Particularly relevant is the agreement with the typically observed right hemisphere inhibitory dominance and the elevated corticospinal excitability in clinical depression [39,43], a disease that includes fatigue as a principal symptom.

The shift in balance to right hemisphere inhibitory dominance could also explain the inability to attend away from current sensory inputs (poor sensory attenuation). Poor sensory attenuation of current proprioceptive and interoceptive inputs, in service of efficient planning of future actions has been proposed as a potential attentional mechanism underpinning post stroke fatigue [12]. Further research is necessary to understand the role of autonomic tone in inter-hemispheric influences and consequently its influence on post-stroke fatigue and other affective disorders.

### Limitations

A potential confounding variable that could compromise the interpretation of inter-hemispheric inhibitory balance is the location of the stroke. To ensure that the side of the stroke did not confound the association between interhemispheric inhibition balance and fatigue and corticospinal excitability, we included the interaction between inhibitory balance and location as a confounding variable. The exclusion of stroke survivors with depression, motoric and cognitive impairments provides a potential bias and a limitation to generalisability of our findings. In this study, stroke survivors with depression were excluded to isolate the group with fatigue without psychiatric comorbidities, as depression has already been associated with aberrant inter-hemispheric connectivity [39,43]. We believe that the exclusion of stroke survivors with depression has not introduced any potential bias to the interpretations of the data, as the focus of the study was understanding biological mechanisms of post-stroke fatigue. We have also excluded participants with large motoric and cognitive impairments, a pragmatic choice that should not lead to limited generalisability of our findings to fatigued stroke survivors with larger impairments. This assumption is supported by a lack of association between IIB and motor and cognitive impairments. The absence of a control group might be viewed as a potential limiting factor – however here we treat fatigue as a continuous dependent variable ranging from minimal to maximal fatigue (1-7), making the interpretation of the findings clinically valid.

### Future directions

The strong explanatory power of resting inter-hemispheric inhibitory dynamics make it a potential target for new intervention protocols using brain stimulation to change corticospinal excitability using transcranial magnetic or electric stimulation. More specifically, brain stimulation methods aiming to recover physiological inter-hemispheric inhibition balance and optimal cortical excitability could be utilised to ameliorate fatigue symptoms [43]. A promising avenue for the future is to combine patient specific characterisation of inter-hemispheric balance, inferred from fMRI or EEG signals, with neuro-stimulation protocols that apply TMS, DC or AC current to rebalance the disturbed inter-hemispheric dynamics. A primary step on this avenue would be to better understand the mechanisms behind the reversal of inter-hemispheric dominance and the relationship between inter-hemispheric balance and neural implementation of attention and sensory attenuation.

## CONCLUSION

We suggest that the balance in inter-hemispheric inhibitory effective connectivity between primary motor regions is involved in regulation of corticomotor excitability and persistence of subjective fatigue. Our findings that post-stroke fatigue and corticospinal excitability are a consequence of altered effective connectivity in M1 support the importance of optimal inter-hemispheric inhibitory balance for healthy brain functioning.

## Acknowledgements

We would like to acknowledge the help of Ms Isobel Turner and Ms Cora Burke during scanning and Dr Ella Clark for her help with the clinical assessments.

